# Active learning’s impact on student course performance in STEM varies by type and intensity

**DOI:** 10.1101/2025.06.01.657285

**Authors:** Shangmou Xu, Vicente Velasco, Mariah J. Hill, Elisa Tran, Sweta Agrawal, E. Nicole Arroyo, Shawn Behling, Nyasha Chambwe, Dianne Laboy Cintrón, Jacob D. Cooper, Gideon Dunster, Jared A. Grummer, Kelly Hennessey, Jennifer Hsiao, Nicole Iranon, Leonard Jones, Hannah Jordt, Marlowe Keller, Melissa Lacey, Caitlin E. Littlefield, Alexander Lowe, Shannon Newman, Vera Okolo, Savannah Olroyd, Brandon R. Peecook, Sarah B. Pickett, David L. Slager, Itzue W. Caviedes-Solis, Kathryn E. Stanchak, Vasudha Sundaravaradan, Camila Valdebenito, Claire R. Williams, Kaitlin Zinsli, Scott Freeman, Elli J. Theobald

## Abstract

We updated a recent meta-analysis of active learning’s impact on student achievement in undergraduate STEM courses by following the same protocol to evaluate studies published from 2010-2017. We screened 1659 papers, coded 1294, and found 210 that met five pre-established inclusion criteria and six pre-established criteria for methodological quality. After further dropping 76 studies with no exam scores data, 134 of these studies contained data on student performance on identical or equivalent exams. We found that on average, active learning’s effect size on exam scores was 0.519 ± 0.049, meaning that when students are in active learning classes, they perform roughly half a standard deviation higher on an identical exam. Funnel plots and sensitivity analyses indicated that these results were not due to sampling bias. Active learning had a positive impact on student outcomes regardless of class size, course level, or STEM discipline, though there was heterogeneity in the effects. All of these results are very similar when compared to earlier meta-analyses, however increased resolution in the studies analyzed here revealed two novel results. First, student performance was significantly better in courses that employed high-intensity active learning, defined as students being on task at least two-thirds of class time, versus lower-intensities. Additionally, there was significant heterogeneity in efficacy across different types of active learning employed. These results suggest that most, if not all types of active learning are effective, and that when innovating in their classes, instructors should continually work to increase active learning intensity. We urge caution in interpreting the results on active learning types, however, and propose a preliminary framework for making more-sophisticated and reliable analyses of variation in course design. Finally, the evidence presented here for active learning’s impact on student outcomes creates a strong foundation for faculty professional development and administration.

## Introduction

A meta-analysis of studies that were published prior to 2010 compared student performance in sections that included any type of active learning to traditional lecture sections of the same undergraduate STEM course and found that on average, exam scores increased by 0.46 standard deviations with active learning and failure rates decreased by 12% (1). Although those and other results inspired urgent calls for converting lecture-focused courses to evidence-based approaches that employ active learning (2), it also inspired passionate defenses of the traditional lecture (3–6).

Recent surveys of undergraduate courses in the STEM disciplines have shown that instructor-centered lecturing still dominates instruction (7,8). For example, data collected with the COPUS classroom observation tool (9) on over 2000 classes given by 500 instructors showed that about 55% of the courses surveyed were interpreted as “didactic” and another 27% as “interactive lecture”—meaning lecture occasionally interrupted by clicker questions or Socratic questioning (8). Even in the 18% of courses classified as “student-centered” in this study, lecturing occurred in at least 50% of the two-minute observation intervals recorded in the COPUS protocol. The researchers also pointed out that they had no data on the quality of the active learning activities observed—only the quantity. This is an important issue, as the quality of active learning can be low due to poor fidelity of implementation for evidence-based approaches (10–13).

Research that contrasts lecturing with active learning is informative because the two broad types of classroom practice are grounded in distinct models of how people learn. First, the *transmissivist theory* of learning describes the instruction approach where the direct transmission of information from an expert in a domain to a novice learner dominates the learning process (14), regardless of the style and the quality of lecturing (3,6). Under this framework, the instructor is the center of a lecture, while students take the role of passive recipients. To reinforce this approach, advocates of lecturing sometimes allude to class sessions as a performance (4). Second, active learning, in contrast, is grounded in *constructivist theory* of learning, which holds that humans learn by actively using new information and experiences to modify their existing models of how the world works (15–18). The “actively using” element in the model is based on the premise that students need to struggle to make sense of new ideas and perspectives, and is reflected in the consensus definition of active learning: Any approach that engages students in the learning process through in-class activities, with an emphasis on higher-order thinking and group work (1,19,20). Therefore, active learning is often characterized as student-centered.

Transmissivist theory vs. constructivist theory is critical to establish distinctive standards for evaluating excellence in college teaching. Transmissivist theory of learning suggests that undergraduates learn best from domain experts, and thus the most-prestigious institutions with the most celebrated domain experts are most capable of providing the best instruction. These domain experts’ job in teaching, in turn, is to synthesize a cutting-edge understanding of the current theory and knowledge in a field and communicate it in the most efficient way possible—the lecture. Under this framework, top researchers who can prepare a well-organized lecture are thought to be the best college teachers, because they have the deepest knowledge of the domain and are actively expanding the discipline’s current frontier. Students act as passive recipients of this distillation and delivery process. In effect, they “learn at the feet of the master.”

Constructivism implies a different standard for excellence in college teaching than transmissivism. Cognitive scientists hold that domain experts often lack knowledge of teaching (21). Under the constructivist model, the best college instructors are not the best domain experts but rather the best experts in teaching a domain. A teaching expert, in this view, has to draw on deep knowledge of the field, but in addition has to understand why students struggle with domain-specific concepts or skills, then design and implement activities that are scaffolded in a way that helps novice learners develop a more-expert-like understanding or skill level. Mary Pat Wenderoth, an early advocate of active learning, characterizes the difference in teaching practice under the transmissivist versus constructivist approach as, “Ask, don’t tell” (22). The skill set required to do this asking successfully starts with domain expertise, but expands to encompass cognitive empathy and social empathy. Cognitive empathy is required to recognize sources of misunderstanding and devise effective solutions; social empathy is required to manage motivation and other aspects of student emotion and psychology—providing support as students struggle to learn.

Moreover, transmissivist and constructivist theories of learning also speak to the development of cognitive skills. Pioneering work by Dweck (23) showed that most individuals adopt one of two perspectives of cognitive skills: A “fixed” mindset that emphasizes innate ability, or a “growth” mindset that emphasizes intellectual growth and the expandability of skills. In an instructor-focused, transmissivist classroom, students undergo a selection process based primarily on aptitude and preparation. In a student-centered, constructivist classroom, the emphasis is on student practice, based on the expectation that all can succeed. Consistent with this claim, recent work has shown that compared with instructors who have a growth mindset, college faculty with a fixed mindset have larger achievement gaps in their classes between racial majority and racial minority students (24), whereas active learning helps reduce such achievement gaps (25).

Constructivism represents a radically different model of learning that contradicts the transmissivism tradition rooted in western-style universities ever since 1088. Because it creates new criteria for pursuing and recognizing excellence in college teaching, it is not surprising that existing faculty have been slow to change. Research has shown that personal empiricism, grounded in the observation that existing faculty thrived under traditional lecturing, is often used to justify teaching decisions and resist change (12). In response, advocates for teaching reform emphasize the need to make evidence-based decisions, parallel to the push for physicians to pursue evidence-based medicine that began in the 1990s (26). In effect, STEM faculty are being asked to use the same evidence-based habits of mind in their teaching as they do in their research (2,27,28).

As such, this study was designed to contribute to the evidence-basis for decision-making in undergraduate STEM instruction by revisiting the Freeman et al. (1) meta-analysis. Specifically, this study’s goals were to update that work by 1. confirming or questioning the effectiveness of implementing active learning with an examination of more-recent empirical work, and 2. exploring the extent to which different practices in student-centered instruction (e.g., different types or intensities of active learning applied) and different classroom settings (e.g., subjects or class sizes) affect student outcomes differently. If accumulating evidence continues to document improved student outcomes in courses that convert from lecturing to active learning, the data support not only a radically different theory of learning, but new standards of excellence in the training and evaluating college faculty.

## Methods

### Overview of the analysis

Following the analytical protocol in Freeman et al. (1), we define traditional lecturing as continuous exposition by the instructor, with student involvement limited to occasional questions, and active learning as any approach that engages students in the learning process through in-class activities, with an emphasis on higher-order thinking (e.g., analysis or synthesis) and group work (1,19,20). The full protocol for this study, presented in the PRISMA-P format (29) and including annotations regarding modifications that occurred during the course of the research, is available in *Supplemental Materials*. *Supplemental Figure S1* provides data on the number of sources that were found and evaluated at each step in the study.

### Literature Search

We searched both gray and peer-reviewed literature for studies conducted or published from January 1, 2010 to June 30, 2016 that reported data on undergraduate performance in the same STEM course under traditional lecturing versus any form or intensity of active learning. We used five methods: hand-searching of journals, database searches, snowballing, mining reviews and bibliographies, and contacting researchers—both selected individuals and the broader community via listserves comprised of educational researchers in each STEM discipline (30–32). See also *Supplemental Table S1* for journals searched by hand for this study.

### Criteria for Admission

The study protocol established criteria for admitting research for potential coding (*Supplemental Materials, Part A*); these criteria were not modified in the course of the study (33). To be considered for coding, sources had to 1. contrast any form and intensity of active learning with traditional lecturing, in the same undergraduate course and at the same institution; 2. involve undergraduates in a regularly scheduled course, 3. focus on interventions that occurred during class time or recitation/discussion sections, 4. treat a course in STEM, which we defined as astronomy, biology, chemistry, computer science, engineering (all fields), environmental science, geology, mathematics, nutrition or food science, physics, psychology, or statistics, and 5. include data on course assessments.

During the original search, papers were downloaded based on information in the title and abstract. One of the co-authors (SF) then checked every resulting paper against the five criteria for admission, based on a check for information in the introduction, methods section, and any figures or tables. In cases that were ambiguous, the paper was referred to coders for a final check on criteria for admission.

### Coding process

Each paper admitted to the coding step was evaluated independently by two of the co-authors. Coders then met to reach consensus on coding decisions for each element in the coding form established in the study protocol (*Supplemental Material, Part A*).

In addition to making a final evaluation on the five criteria for admission, coders had to reach consensus on the data to use in computing effect sizes and the data used in moderator analyses, including the STEM discipline, course level (introductory versus upper-division), intensity of the active learning intervention in terms of the percentage of class time devoted to student-centered activities (*Supplemental Table S2*), and the type of active learning involved (*Supplemental Table S3*).

Coders also evaluated information designed to minimize both within-study bias and across-study bias (29,34):

i. To control for the impact of time on task on performance, we excluded studies where class time per week was longer in the active learning treatment.
ii. To control for the impact of class size on performance, average class size could not differ by more than 25% of the larger class, unless the active learning section was larger.
iii. To control for examination equivalence, the assessment coded as an outcome variable had to be identical, formally equivalent as judged by an independent analysis performed by experts who were blind to the hypothesis being tested, or made up of questions drawn at random from a common test bank.
iv. To control for student equivalence, students either had to be assigned to treatments at random or, for quasi-random studies, reports had to include data indicating that there was no statistically significant difference between students in each treatment group in terms of entrance exam scores, high school or college grade point averages, content-relevant pre-tests, or other measures of academic preparedness and ability.
v. To assess the impact of instructor ability on performance, the instructors in the treatments were coded as identical, drawn at random from a pool, or comprised of a group of four or more in each case—meaning that both treatments were team-taught.
vi. To avoid pseudoreplication when a single study reported the results of multiple experiments—for example, how studio course designs impacted student performance in more than one course or institutional context—the coders identified comparisons of treatment and control conditions that were independent in terms of both course identity and student population. Data from multiple sections of lecturing and/or active learning in the same course and institution were recorded separately, but were meta-analyzed as a single study.

If studies met both the admission criteria and the six quality criteria just listed, but did not include all of the data required for admission to the study, we contacted all authors to request the missing data.

As a final quality control step, one of the co-authors (MJH) checked all studies and all codes against the source information to resolve any inconsistencies or ambiguities, consulting with a second co-author (ETran) to resolve any uncertainty. *Supplemental Table S7* summarizes the list of studies and active learning features of each selected study.

### Analytical approach

The dataset used in the final analysis contained 37,839 student records from 133 independent studies with data from identical or formally equivalent exams. Each study represented a specific course and active learning intervention; the data came from 8 different STEM disciplines and included both introductory and upper-level courses, several active learning types, taught at three intensities (*Supplemental Table S4*). Of the 134 studies, 90 reported results from quasi-random experimental designs, as students self-sorted into classes with different treatments (15 were random and the design of 29 were unknown).

Many studies reported data from multiple active learning sections and multiple traditional lecture sections, with no clear pairing. Thus, to avoid unnecessary multiple comparisons, we took a weighted average of the exam scores and their standard errors in the traditional lecture sections and using that average, we then computed the standardized mean difference and standard errors for each active learning section of the same study (1). This gave us a single pairwise comparison for each study with an exception when a study reported multiple comparison results. In such cases, we extracted multiple comparisons from a single study; we term each of these comparisons “case studies.” In the 133 studies in the meta-analysis, we have 232 case studies. When multiple case studies were retrieved from single studies, we computed multiple estimates (one per case study), thus estimates from these single studies were not independent. We accounted for this non-independence in two ways. First, we included a study random effect which accounts for the fact that the errors are correlated. Second, because the same control group (albeit a weighted average) was used to calculate multiple effects, we estimated the variance-covariance matrix of the effect size matrix prior to fitting the models, following recommendations from Gleser and Olkin (35).

In addition, because this meta-analysis investigates student performance and students are nested within classrooms, there is also non-independence between the exam scores of each student. It was rare in our dataset for the original authors to account for this non-independence (for example, with a random effect) (36). Thus, to account for the non-independence between students, we used cluster correction methods developed by Hedges (37). This cluster correction relies on the intraclass-correlation (ICC), which is used to measure the relationship between the variability within each cluster (classroom) and across clusters (classrooms). We expect the variability across clusters to vary, depending on the level at which the measure is made. For example, if the ultimate comparison is between two sections within the same term there should be more non-independence than if the comparison is between two iterations of the class in different years. To account for this difference in variability, we used various ICCs, clustering by Section, Term, or Year, depending on the level at which the comparison between groups of students was made in the original publication. We obtained an empirical estimate of these ICCs by considering more than 25,000 chemistry students clustered within 6 classes in 4 quarters per year over the course of 16 years at a single university (data from Harris et al. 2020) (38). Thus, we defined our ICC to be 0.093, 0.002, and 0.003 for Section, Term, and Year, respectively.

We calculated the effect sizes as weighted standardized mean differences (Hedge’s g) to analyze the effectiveness of the different factors of active learning compared to traditional classrooms. We fit multilevel models using the rma.mv function in the *metafor* package (39). All analyses were conducted in R version 4.0.5 (40).

### Bias Assessment and Quality Assurance

To reassure ourselves that our results are not heavily influenced by publication bias we performed various methods of Bias Assessment and Quality Assurance. First, to test whether the small studies in our sample are skewed by publication bias we created funnel plots and visually assessed the symmetry. While this is recommended with meta-analyses that do not include random effects, this is not a method that is highly recommended given our model fitting procedure. Specifically, our models included multiple effects from the same study, thus many of the points in the funnel plot are not considered independent from each other (41). Therefore, true asymmetry is difficult to assess in our sample. Nevertheless, there is no obvious asymmetry in the funnel plots (*Supplemental Figure S2*).

Next, we calculated Failsafe-N values with both the Rosenthal (42) and Orwin (43) approach. The Rosenthal approach considers the hypothetical of how many studies with zero effect would be needed to take the *p*-value of the grand mean model over 0.05. The Orwin method takes a slightly different approach and instead looks at effect size, estimating the number of studies with zero effect that would be needed to reduce our calculated effect size to half of the estimated effect size. From the Rosenthal approach we found that 29,994 studies with null effects would need to be added to our data set to make the *p*-value of our results non-significant (i.e. > 0.05). From the Orwin approach, we found that we would need to add 232 papers with an effect size of zero to cut our calculated effect size in half.

Additionally, we plotted the distribution of *z*-scores of all the estimates in our meta-analysis. We did this to observe the amount of studies in our sample that had *z*-scores around the conventional threshold of statistical significance (z = 1.96, p = 0.05). If our meta-analysis was influenced by publication bias we would see a dip in the amount of studies in our sample right below the threshold (z < 1.96, p > 0.05), meaning that there are likely studies that were non-significant which were not favored by publishers and not included in our sample (44). Similarly, if there was a spike in studies just past the threshold (z > 1.96, p <= 0.05) this would indicate that publishers were favoring studies with statistically significant results leading to our data set being subjected to publication bias. We see that there are no spikes past the threshold and no dips before the threshold providing more evidence that our results were not influenced by publication bias (*Supplemental Figure S3*). Through all these tests, we conclude that there is little to no evidence of publication bias in our data set.

Another form of bias would be if our coefficient of non-independence (Intraclass correlation coefficient, ICC, or rho) does not accurately represent the non-independence of the students in the classes and the classes in the studies in our synthesis. We estimated the rho values we used empirically from a different dataset (see Methods) but to assess whether our established rho heavily influences the results we report, we recalculated our grand mean effect size using a variety of rhos (0.093, 0.1, 0.22). *Supplemental Table S5* shows the estimates as well as the standard errors of these models estimated with different rho values. For no value of rho does the grand mean effect size change such that the effect is statistically indistinguishable from zero, thus our selection of rho does not heavily influence our results. We choose to interpret the model that uses rho = 0.093, 0.002, and 0.003 for Section, Term, and Year, respectively because that is the rho value that most closely aligns with the rho values calculated in a large dataset of chemistry classes at a large university (38).

Finally, to test whether there are single studies which contribute disproportionately to the overall results of the meta-analysis we conducted an undo-impact analysis. Here, we individually removed a single study and recalculated the grand mean effect size. *Supplemental Figure S4* shows a histogram of the grand mean effect sizes from this analysis. In every iteration of refitting the model, the grand mean effect size always stayed significantly positive. This indicates that no single study had an undo impact on the results we report here. Relatedly, we also consider whether an alternative form of random effects may change the estimation of the effect sizes. The final model only includes a random effect for section, not for author. As indicated in *Supplemental Table S6*, no author had an extremely large or small standard deviation of the residuals (mean=0.9661, sd=0.046). All mean residuals (by author) were close to zero (mean=0.0065, sd=0.0266). This indicates that accounting for non-independence within section is sufficient, and a within-author random effect is not necessary.

## Results

We present the estimated effectiveness of active learning on students’ performance compared to traditional classrooms by visualizing effect sizes (Hedge’s *g*) and the associated standard errors. *Figure 1* summarizes the overall effect of active learning and different factors that may influence the effectiveness of active learning. The top panel of Figure 1 examines the overall effect of active learning, whereas the next two panels concern the extent to which different practices of active learning may change the effect sizes. Finally, the bottom three panels of Figure 1 examine the influences of classroom settings on the effectiveness of active learning.

**Figure 1.**
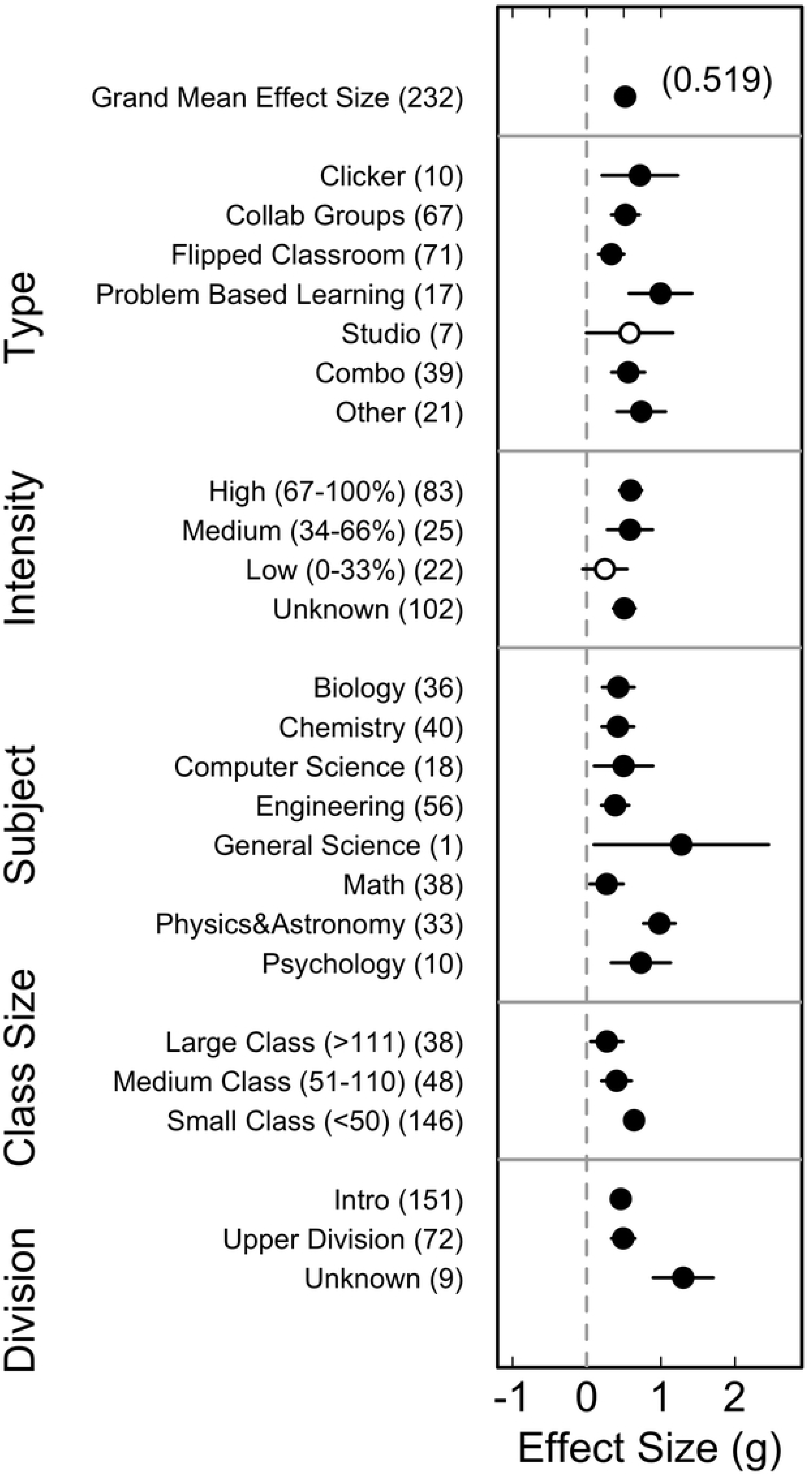
Visual summary of the effectiveness of active learning by practices and classroom settings (n = 232 case studies).

The top panel in Figure 1 reports the average effect size of active learning on students’ performance by pooling all different types of classrooms. The effect size is captured by weighted mean differences of performance between traditional instructor-centered classrooms and classrooms with active learning. The estimated overall effect size is 0.519 (*Z* = 10.537, *p* < 0.001, *df* = 231), indicating that, on average, students in active learning classes performed approximately half a standard deviation higher than their counterparts in traditional lecturing. To help visualize the variation of the individual effect sizes across all selected studies, *Figure 2* depicts a caterpillar plot that indicates the distribution of the effect sizes. As shown in Figure 2, most effect sizes are within the range from 0 to 2, with some extreme sizes at both sides.

**Figure 2.**
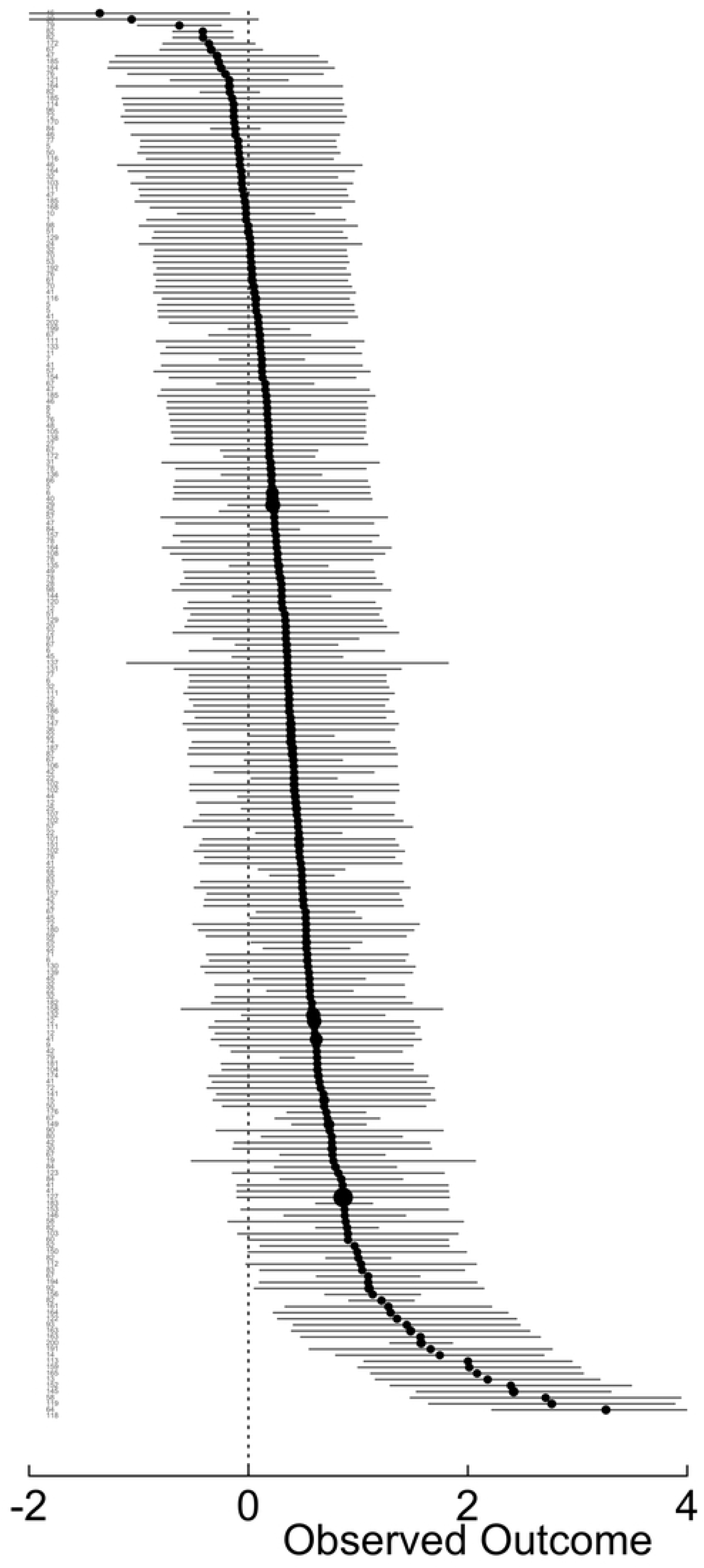
Caterpillar plot of the mean effect sizes across all selected studies.

We then implemented subgroup heterogeneity analyses to understand the extent to which active learning practices and classroom settings, as moderators, influenced the effectiveness of active learning. As visualized in Figure 1, overall, there were noteworthy differences in effects of active learning across different contexts, including intensity and type of active learning and classroom settings. *Table 1* further summarizes the heterogeneity analyses with estimated effect sizes of each subgroup and their corresponding Cochran’s Q and *I*^2^ statistics. In general, most *I*^2^statistics reported in Table 1 were approximately at a 0.6 level, indicating that 60% of the observed variance was due to the differences across subgroups. Particularly to different subgroups, as shown in Table 1, Panel 1, almost all types of active learning (except studio, likely due to low sample size) had a positive effect on students’ performance, with substantial variability across different types of active learning implemented (*Q*_*M*_ = 120.5, *df* = 7, *p* < 0.001; *I*^2^ = 59.8%). Similarly, Panel 2 shows that there were statistically significant differences in the impacts on exam scores across different intensity levels of active learning (*Q*_*M*_ = 112.8, *df* = 4, *p* < 0.001; *I*^2^ = 61.1%), with middle and high intensity of active learning exhibiting significantly positive effect sizes. However, low intensity active learning did not have a significantly different impact on exam scores than lecturing.

**Table 1.**
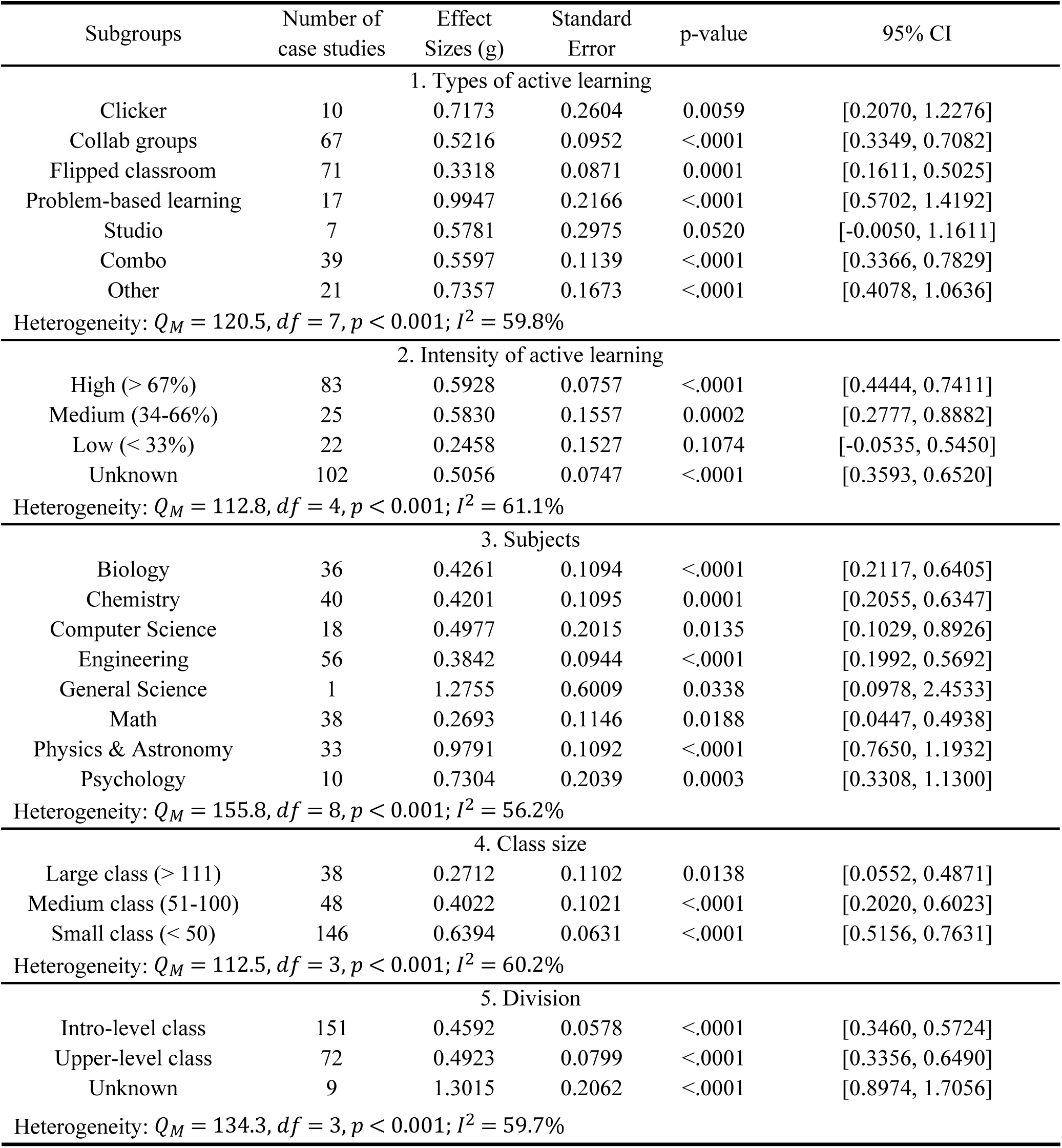
Summary of sub-group heterogeneity analysis: Moderator effects of different factors including types and intensity of active learning, and classroom settings.

There were differences in the impact of active learning across different classroom settings: Panel 3 in Figure 1 shows that there were differences in effects of active learning across different disciplines (*Q*_*M*_ = 155.8, *df* = 8, *p* < 0.001; *I*^2^ = 56.2%), but we found that active learning had a significantly positive effect on exam scores in every discipline. Additionally, we found that active learning had a significant positive effect on classroom exam performance regardless of the class size (Panel 4). Yet, there was a slightly larger effect size, on average, in smaller classes (*Q*_*M*_ = 112.5, *df* = 3, *p* < 0.001; *I*^2^ = 60.2%), indicating that the effectiveness of active learning is amplified in smaller learning environments. Finally, we found that in both introductory and upper-level classrooms, active learning had significant positive effects on students’ performance (Panel 5). Yet, students in upper division courses benefited marginally more from active learning than students in introductory courses (*Q*_*M*_ = 134.3, *df* = 3, *p* < 0.001; *I*^2^ = 59.7%).

## Discussion

Earlier meta-analyses have shown that all students benefit from active learning in undergraduate STEM courses compared to traditional lecturing (1,45,46). The results of those earlier studies are consistent with the data reported here, documenting an improvement in exam scores of about half a standard deviation, averaged over courses with different active learning practices and classroom settings in undergraduate STEM education. Furthermore, our results extend these results to interrogate the impact of active learning, given the type and intensity of active learning and other classroom settings. Perhaps most importantly to educational practitioners, in most contexts, active learning exhibits positive effects on student performance. To name a few, active learning can improve student outcomes regardless of types of active learning techniques, subjects, and class division. Yet, we also found that active learning exhibits greater impacts on performance in smaller classrooms and when used for more than a third of in-person instructional time. These insights have important implications for teaching practice and for discipline-based education research.

### Implications for teaching practice

The results reported in this paper increase confidence that, on average, active learning, a constructivist approach, in undergraduate STEM courses results in greater learning outcomes compared to a “transmissivist” lecturing approach. As a result, our data support calls for instructors to convert their teaching practice to an evidence-based framework (2)—especially in light of recent surveys documenting that current practice in undergraduate STEM courses is still dominated by lecturing (8,47).

The results reported here highlight a formidable challenge to faculty change, however, based on the role of active learning intensity in explaining variation in student outcomes. We call this challenge the “start-small conundrum.” Most experts in faculty development urge instructors who are new to active learning to make a few initial changes to their classroom practice before gradually increasing the percentage of class time devoted to student-centered teaching. This recommendation is motivated by two observations: 1. Instructors can rarely, if ever, completely redesign a course before teaching it again while also fulfilling their other professional obligations, and 2. initial efforts often fail. Early failure is common—even expected—because instructors have to learn how to design effective materials and manage an active classroom successfully, in addition to customizing their curriculum and classroom culture to their student population (48). Unfortunately, researchers have documented that one third of faculty who try active learning approaches abandon them after an initial attempt (27). If faculty who revert to lecturing claim that “active learning didn’t work,” the data reported here support their rationale if the initial implementation lacked sufficient intensity. In practice, educators may overhaul their teaching philosophy and invest hundreds of hours in redesign—yet observe no measurable gains in student performance unless they reach a threshold of engagement time.

Overcoming this “start-small” barrier requires both individual persistence and institutional support. Persistence, or Grit, is defined as sustained passion and perseverance toward long-term objectives (48,49) that enables instructors to iterate on course design past an initial setback. Figure 1 demonstrates that, on average, instructors who continue modifying their course and ultimately meet the high-intensity time-on-task criteria, or reach “collaborative learning” level described in Lund et al. (47), achieve statistically significant improvements in student outcomes.

One earlier meta-analysis explicitly evaluated the impact of active learning intensity on student outcomes (25), and extensive earlier literature nonetheless emphasizes the importance of time on tasks. For example, Freeman et al. (1) analyzed nine outliers with particularly large and positive effect sizes in their sample, and found that seven were studio course designs or other approaches where essentially 100% of class time is devoted to active learning. In addition, a series of reports in the literature have shown a strong positive relationship with student outcomes— usually exam or concept inventory scores—when the percentage of class time devoted to active learning was a key variable in the study design (10,50–55). In one particularly careful study, Weir et al. (55) estimated that a 10% increase in group work time in class was correlated with an increase of 0.3 in the effect size on student outcomes—in this case, about a 3% improvement in pre-post comparisons of concept inventory gains. Similarly, a recent meta-analysis of active learning’s impact on low-income and underrepresented minority students showed that achievement gaps in both exam scores and failure rates only closed in course designs where more than 66% of class time was spent in active learning (25). Finally, the literature on deliberate practice emphasizes the importance of extensive and strenuous effort to achieve improvement on both academic and non-academic tasks (56). Taken together, the data reported here and in the literature suggest that more is better when it comes to active learning—although quality, defined as fidelity to evidence-based implementation (25), is essential (10,57).

Finally, we urge caution in interpreting our observation of heterogeneity in student outcomes based on the type of active learning employed. Although we established objective criteria for classifying interventions (*Supplemental Table S3*), most published papers lack the granular implementation details that other instructors could repeat the intervention and expect the same outcomes. Therefore, we do not recommend that practitioners preferentially adopt one format (e.g., clicker questions or problem-based learning) over another (e.g., flipped classroom or groupwork), despite the fact that these approaches exhibited significantly different mean effect sizes in our analysis. Instead, instructors should focus on meeting the high-intensity, time-on-task thresholds shown here, in combination with high-fidelity implementation, to drive consistent gains in student learning.

### Implications for research

Calls for reform in undergraduate STEM education began over 25 years ago (28,58,59), and evidence on the efficacy of active learning, including this study, has been accumulating ever since (1,25,45,46). These observations raise a key question for discipline-based education research: Why have STEM faculty been so slow to change their teaching practice? The answers emerging from research cite an array of impediments, ranging from a reliance on personal experience instead of research syntheses, faculty identifying as researchers rather than teachers, a lack of rewards for successful innovation, and an over-reliance on top-down mandates from administrators or bodies of national experts versus bottom-up, community-based support (11,12,60). The novel insight on the role of active learning intensity reported here suggests that if researchers alert faculty to the start-small conundrum and normalize early failure, paths to innovation might become slightly more tractable.

Our results also support calls for discipline-based education researchers (DBER) to abandon experimental designs focused on contrasting active learning and traditional lecturing, moving instead to what observers have called 2^nd^-generation education research or Education Research 2.0 (1,2,61). Instead of asking whether active learning leads to better student outcomes on average, the relevant questions now focus on why active learning works and how to design interventions that benefit specific student groups or teach particularly difficult concepts and skills. Relatedly, this study speaks to recent critique (e.g., Martella et al., 2023) of lack of internal validity in active learning research (62). Our results indicate that active learning can improve student learning outcomes regardless of the choice of active learning practices and classroom settings, providing some evidence that the observed effects of active learning are less likely due to differences in background such as classroom settings.

In terms of guiding the direction of 2^nd^ generation education research, we recommend that researchers refrain from comparing “types” of active learning strategies and instead focus on changing specific elements of active learning practice or other aspects of course design. To support this change, the DBER researchers on course interventions, and the editors and reviewers who evaluate them, should apply the same standard for reporting methods that is expected in other STEM fields: Based on the information in the paper, a qualified individual in a different research group should be able to replicate the study. To meet this standard, discipline-based education researchers should refrain from reporting interventions primarily on the basis of named types such as flipping, problem-based learning, clickers, or studio. Instead, we propose that experimental designs should quantify as many aspects of the intervention as possible, focusing on those that are known to impact student learning (63).

Specifically, we propose that intervention studies are not considered publishable unless experimental and control treatments are quantified with one or more validated classroom observation tools (e.g., PORTAAL or BERI). Multiple tools may be required in some studies, as some of the published instruments focus exclusively on quantifying aspects of student engagement or student-instructor interactions (64–66), while others quantify classroom activity (9,67–72).

In addition to documenting classroom practice during face-to-face sessions, published reports on active learning interventions should include data on assigned work before class sessions, assigned work after class sessions, and other aspects of course design, such as alignment between formative and summative assessments and learning goals and objectives. Although several published instruments implement at least some interventions (73–76), researchers may need to verify the student- or instructor-reported data in these tools with independent assessments.

Our call for researchers to transition from relying on named course design strategies to quantifying actual practice is timely: Most observation tools cited earlier were published during or after the 2010-2016 interval analyzed in this study. Stated another way, the tools required to quantify practice and make interventions more repeatable were published during this period. Given the breadth and quality of instruments now available, however, it is now contingent upon the community to mandate their use as a standard in publishing discipline-based education research.

### Implications for faculty training and evaluation

The evidence for active learning’s impact on student outcomes creates a strong ethical foundation for innovation in undergraduate instruction (2). Recent results showing that high-intensity active learning reduces the inequities in student outcomes that are observed under traditional lecturing— specifically, the disparities experienced by minoritized students (25)—adds a social justice imperative to the ethical element driving faculty change. The research reported here and elsewhere presents a strong case for making constructivist theory and evidence-based teaching central to faculty training and evaluation.

For mentors tasked with preparing future faculty, the constructivist model means that becoming a well-trained and highly accomplished domain expert is, by itself, insufficient as a training and hiring criterion for college instructors. Instead, graduate and post-doctoral advisors and program managers need to support mentees who are interested in faculty positions by offering seminars that introduce the literature on evidence-based teaching and apprenticeships with faculty who are experienced with implementing active learning.

For administrators tasked with improving the teaching skills of existing instructors, the constructivist model means that faculty development becomes an urgent priority. Specifically, professors will need to be supported, through faculty learning communities or other structures, as they transition from lecturing to active learning—especially during the early, low-intensity phases when they are unlikely to observe better student outcomes. Even more important, efforts to adopt evidence-based course designs and eventual success with improving student outcomes will need to be rewarded during merit pay, promotion, and tenure decisions.

Based on the data presented here and elsewhere, we urge administrators to recommend that faculty submit data from classroom observation tools, lists of learning objectives, samples of pre-class preparation materials and post-class exam practice opportunities, and evidence of alignment across learning objectives, assessments, and teaching practice as part of their merit pay proposals and promotion and tenure packages. Awards and other forms of financial and professional recognition should require data on improved student outcomes. These recommendations are timely, as a number of institutions and professional societies are exploring reforms to traditional student evaluations of teaching (SETs) (77). This reform movement, in turn, is a response to data documenting both gender and other biases in SETs and a lack of correlation between SETs and actual student learning (78,79).

### Limitations

Much like many meta-analyses, this study seeks to summarize the average impact of two treatments over many populations, local circumstances, and details of implementation. Here our goal was to summarize the average impact of active learning compared to lecturing over many student populations, STEM courses, institution types, and instructors. In addition, our decision to dichotomize lecturing and active learning masks substantial variation in both practices. In interpreting the results reported here, then, it is important to recognize that they are only valid on a summary, “on-average” level. Although we followed best practices designed to limit the influence of the file drawer effect and other sources of bias within and between studies, no meta-analysis can claim to be free of bias. Confidence in our results rests on their consistency with conclusions from previous studies (1) and assessing how well our methods align with best practices.

## Conclusion

“Transmissivism” is a theory of learning that predicts better student outcomes when highly accomplished domain experts present novice learners with clear, well-organized expositions summarizing the salient models, knowledge, and skills in their fields. Constructivism is a theory of learning that predicts better student outcomes when novice learners are required to actively engage with new problems, models, and data. In testing these contrasting predictions, the data reported here and elsewhere support constructivist theory and reject the transmissivist model in undergraduate STEM education.

## ACKNOWLEDGMENTS.

We thank Darrin Howell for help with hand-searching journals. Financial support was provided by the University of Washington, Office of the Dean of Arts and Sciences.

